# Link Your Sites (LYS) Scripts: Automated search of protein structures and mapping of sites under positive selection detected by PAML

**DOI:** 10.1101/540229

**Authors:** Lys Sanz Moreta, Rute R. da Fonseca

**Affiliations:** The Bioinformatics Centre, Department of Biology, University of Copenhagen, Denmark; Center for Macroecology, Evolution, and Climate, Natural History Museum of Denmark, University of Copenhagen, Copenhagen, Denmark

**Author notes:** Correspondence: Lys Sanz Moreta, / /.

**Keywords:** Functional domain, positive selection, BLAST, PDB, codeml, homologous proteins, Prosites, pymol

## Abstract

The visualization of the molecular context of an amino acid mutation in a protein structure is crucial for the assessment of its functional impact and to understand its evolutionary implications. Currently, searches for fast evolving amino acid positions using codon substitution models like those implemented in PAML [1] are done in almost complete proteomes, generating large numbers of candidate proteins that require individual structural analyses. Here we present two python wrapper scripts as the package *Link Your Sites* (LYS). The first one i) mines the RCSB database [10] using the blast alignment tool to find the best matching homologous sequences, ii) fetches their domain positions by using Prosites [3,8,9], iii) parses the output of PAML extracting the positional information of fast-evolving sites and transform them into the coordinate system of the protein structure, iv) outputs a file per gene with the positions correlations to its homologous sequence. The second script uses the output of the first one to generate the protein’s graphical assessment. LYS can therefore generate figures to be used in publication highlighting the positively selected sites mapped on regions that are known to have functional relevance and/or be used to reduce the number of targets that will be further analyzed by providing a list of those for which structural information can be retrieved.

**Motivation:** Automatizing the search for protein structures to assess the functional impact of sites found to be under positive selection by codeml, implemented in PAML [1]. Building publication-quality figures highlighting the sites on a protein structure model that are within and outside functional domains. reduces the workload associated with selecting proteins for which a functional assessment of the impact of mutations can be done using a protein structure. This is especially relevant when analyzing almost complete proteomes which is the case of large comparative genomic studies.

**Software:** LYS scripts are executed in the command line. They automatically search for homologous proteins at the RSCB database [10], determine the functional domain locations and correlate the positions pointed by the M8 model [1], and output a data frame that can be used as the input by PyMOL [7] to generate a visualization of the results.

**Availability:** LYS is easy to install and implement and they are available at https://github.com/LysSanzMoreta/LYSAutomaticSearch

## 1. INTRODUCTION

One of the goals in comparative genomics studies is to find regions of the genomes that evolve at elevated rates, which can potentially indicate that they involved in promoting adaptation to new environments. Such regions are said to be evolving under positive selection [10]. It is possible to infer positive selection occurring in individual protein sequences by assessing the rates of substitutions at specific codons (sets of three nucleotides that correspond to an amino acid) thanks to site models such as those implemented in PAML [1,5].

Positive selection is evaluated through the ω value that corresponds to the ratio between the amount of non-synonymous mutations per non-synonymous site and the amount of synonymous mutations per synonymous site. Non-synonymous mutations can be relevant if the amino acid switch introduced generates a change in the physicochemical properties of the residue and consequently affects the protein function. A first step in the evaluation of the impact of these mutations consists on identifying their location on a protein structure (which could be the structure of a closely related homologous protein) and verify whether they are located within known functional domains. In a protein structure the amino acids form a backbone that is folded into a specific conformation, with the folding patterns being dictated by a series of non-covalent bonds (hydrogen bonds, ionic bonds and van der Waals attractions) directed by the residue’s side chains. If the residues in the functional domain are exchanged with an amino acid with different properties, these interactions will be modified together with the structure and its binding attributes will be affected [2]. Mutations in the functional domain are more likely to affect the protein’s function when compared to those located in other parts of the structure.

In order to easily assess which proteins in a large selection scan can be analyzed at the structural level, we present a python wrapper that reads a file containing the sequences to analyze and the paths to the output files from M8 codeml model, and performs an automatic search of homologous proteins by blasting the query sequences to the RCSB database [10]. The results from blast are ranked according to the percentage of identity, the coverage and finally, resolution of the crystallographic protein information file. The selected PDB files are further analyzed via the Prosites [8,9] software implemented in biopython [3] to find the domains positions. Next, the positions correspondence algorithm is implemented among the query sequence and the homologous protein sequence. This correspondence is outputted as a data frame that is then used in a second script to create the visualization of the protein structure with highlighted functional domains and positively selected sites in PyMOL [7].

## 3. METHODS

### Design of the algorithm to perform the positions correspondence

The main algorithm finds the correspondent positions among the query gene sequence and the crystallography file sequences. These are the main steps (see also Figure 1) followed in the script:

1. Creation of two lists: i) list A containing the positions in the alignment (biopython’s [3] global alignment) where there are no gaps in any of the sequences and ii) list B, which has the length of the gene sequence, filled with ‘nan’ values.
2. Counting the amount of gaps between each segment, bounded by the *i* and *i +1* positions contained in list A, in the aligned sequences. This step is performed for both sequences. Two output lists are generated (C and D) with the reciprocal correspondence of the positions where there are not gaps in the alignment of chain A and B.
3. Lists C and D are used to fill in list B with the correspondent positions. Furthermore, the correspondent positions of the gene in the PDB sequence are substituted by the actual residue ID numbers from the PDB file, which follow their own numbering settings.

**Figure 1.**
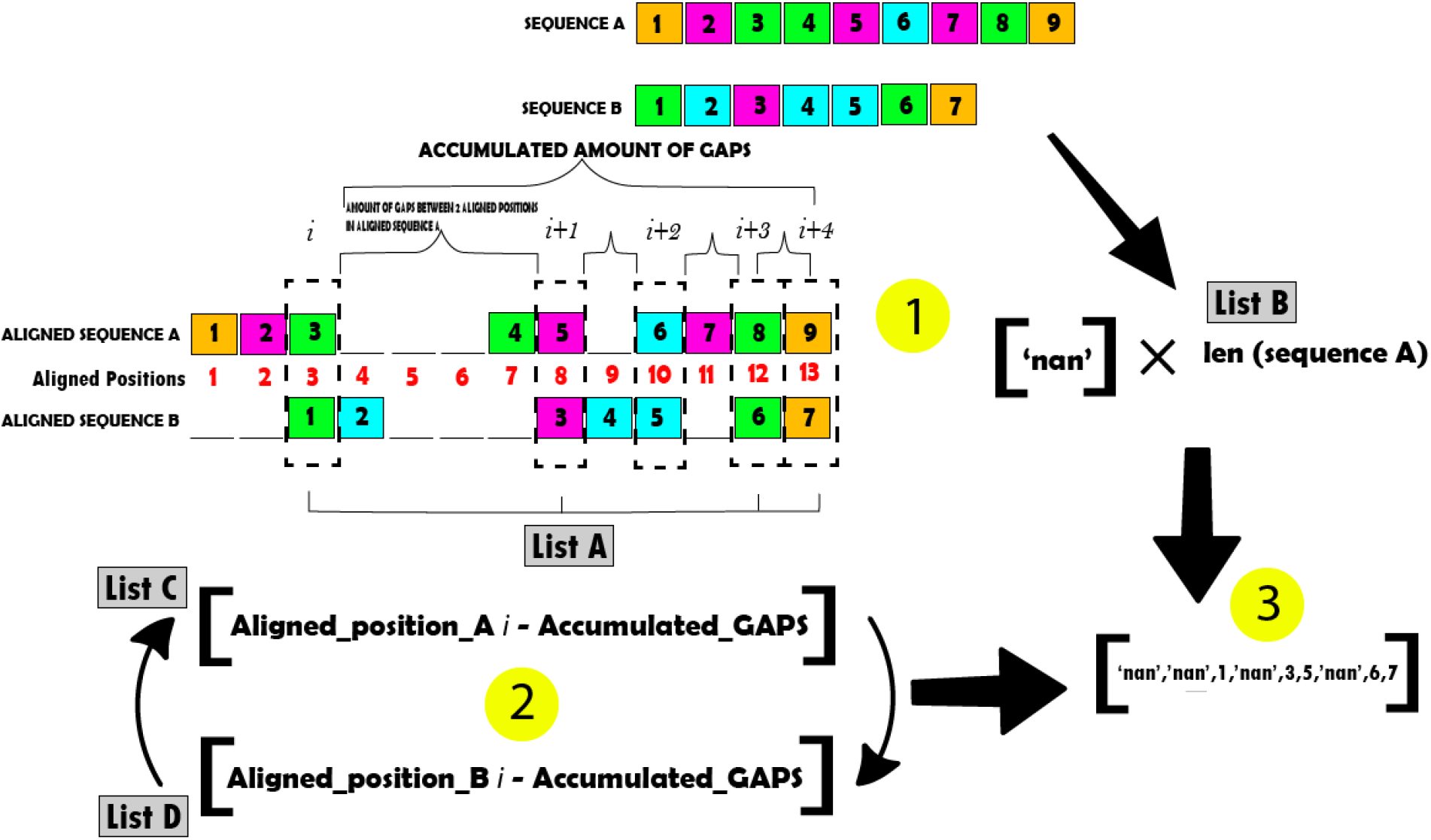
Graphical explanation of the algorithm that matches the coordinates of 2 sequences by using their unaligned and aligned versions (local or global alignment in biopython, 2009). The numbers indicate the residues positions in the chain/sequence.

## 4. MATERIALS

LYS consists of a series of Python version 3 scripts available in a Github repository (https://github.com/LysSanzMoreta/LYS_Automatic_Search) and licensed under an Apache Version 2 License. All of the scripts require the freely available packages of pandas, numpy, pymol and biopython, whose installation is highly recommended through anaconda version 3. The scripts that call Pymol [7] can be also used freely under educational purposes. A simple video tutorial for the two main scripts is available at https://www.youtube.com/watch?v=8ui1TxpOd6M.

LYS has been tested on Unix platforms like Ubuntu 16.04 and Linux. To be able to make use of the scripts that call the pymol GUI, make sure that the pymol Educational version is the in the command line path. The input files for the main script LYS_PDB_Search.py are, a file containing all the sequences (whose formats can be specified with the flag –format, fasta is default and recommended) and a tab separated file containing rows with the name of the sequence (containing the exact same sequence name as in the first file, for example the Fasta headers) and its path to the codeml M8 output results. The complete list of available arguments is shown in Table 1. The outputs, which will be stored in the *Positions_Dataframe* folder, are dataframes containing the positions number correspondence among the input sequence and its selected homologous sequences as seen in Table 3 (currently only the top scoring 3 RSCB crystallography files sequences are chosen, it can be easily changed inside the script). Alongside a folder where the crystallography protein files are downloaded is created (*PDB_files*).

**Table 1.** LYS_PDB_Search.py script list of arguments.

| Argument | Required | Help | Default value |
| --- | --- | --- | --- |
| --Proteins | True | Path to File containing the coding sequences (Recommended Fasta format) |  |
| --Codeml | True | Path to file containing rows with: "Gene name" + '\t' + "Path to codeml M8/bsA1 output file". Remember: Gene name needs to match the Gene name in the Sequences file |  |
| --format | False | Sequence or Multiple Alignment File Format | fasta |
| --prob | False | Choice of level of posterior probability on the sites, 95% or 99% from M8 Out file | 99% |
| --missing_data | False | Decide if the missing data (labeled as "N") should be kept from the nucleotide sequence. It might affect the final alignment, is recommended to check the alignment scores in both options (activate print_alignment to do so). | yes |
| --print_alignment | False | Choose to visualize the PDB file sequence aligned with the gene | no |

Once the data frames have been created navigate to that folder and find, for example through *grep –rl “Selected_and_Domain”*, which ones have determined that the homologous protein displays positively selected residues in the domain. Following, call the LYS_PyMOL_input_Dataframe.py GUI interface, Figure 2, to plot in a personalized approach the proteins, check for customizable features in Table 2, that display the result of interest, Figure 3.The list of available scripts is the following:

**Figure 2.**
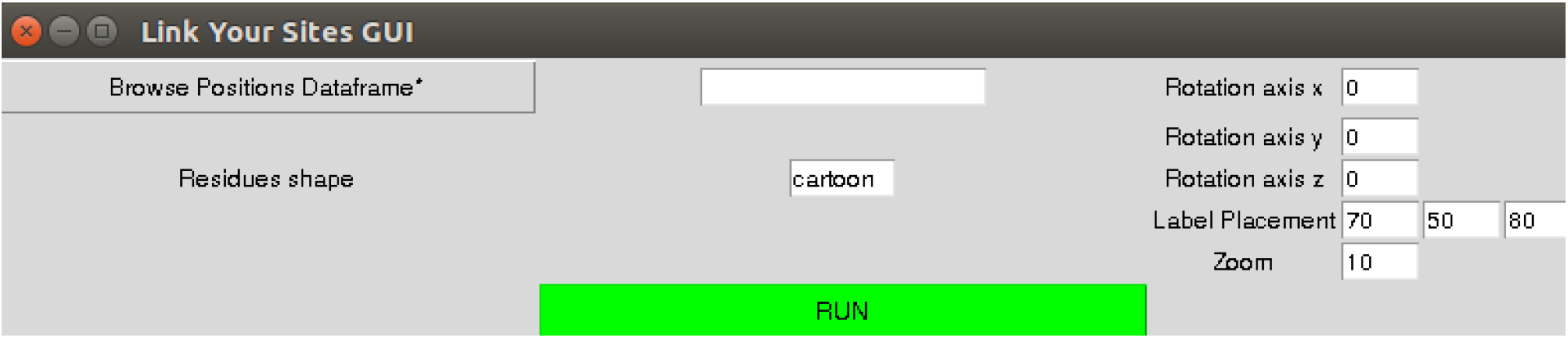
LYS_PyMOL_input_Dataframe.py’s interface. The compulsory files for the GUI to work are marked with a *. Tutorial at https://www.youtube.com/watch?v=8ui1TxpOd6M

**Figure 3.**
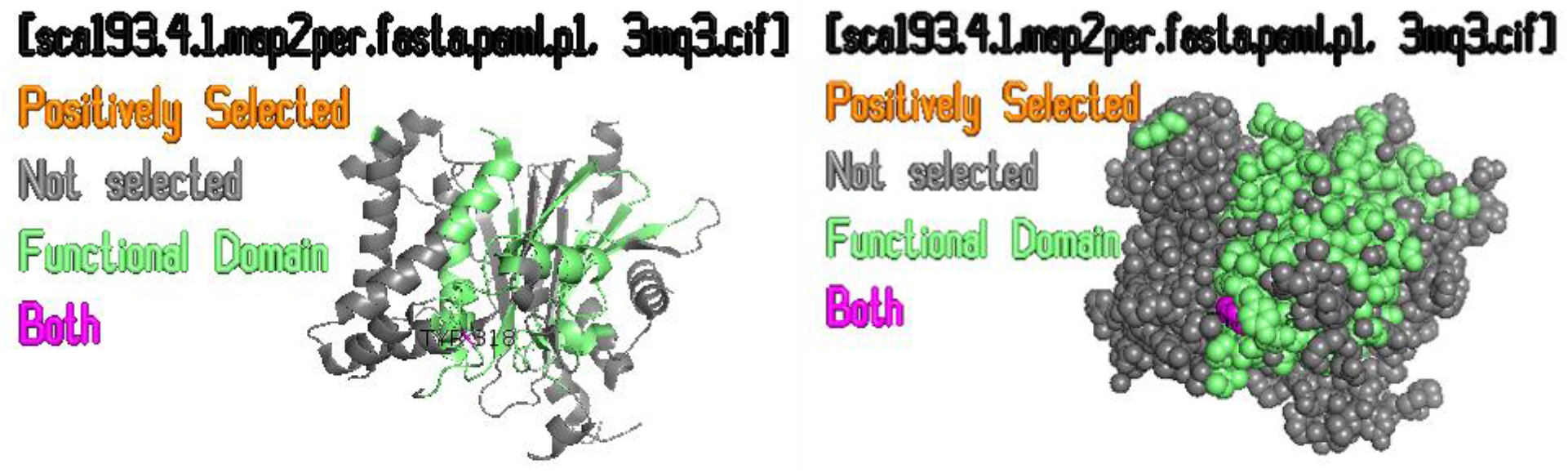
LYS’s visual output examples of protein coloured according to its evolutionary positively selected amino acid residues and domain positions. Cartoon(left) and Spheres(right) modes.

**Table 2.** LYS_PyMOL_input_Dataframe.py customizable features inside the script or GUI.

| Settings | Options |
| --- | --- |
| Background, Residues and Font Colours | Choose colours from the palette:<br>https://pymolwiki.org/index.php/Color_Values. |
| Residues Shapes (GUI) | Choose from:<br>https://pymolwiki.org/index.php/Show |
| Select and Remove chains | Chose if any of the chains should be removed in the visualization. |
| Legend : Font Size and Placement (GUI) | Change the values of the axes, cyl_text and cmd.set |
| Carbon-alpha residues labelling | Activate cmd.label accordingly: Designed to highlight only alpha carbons of selected sites |

**Table 3.**
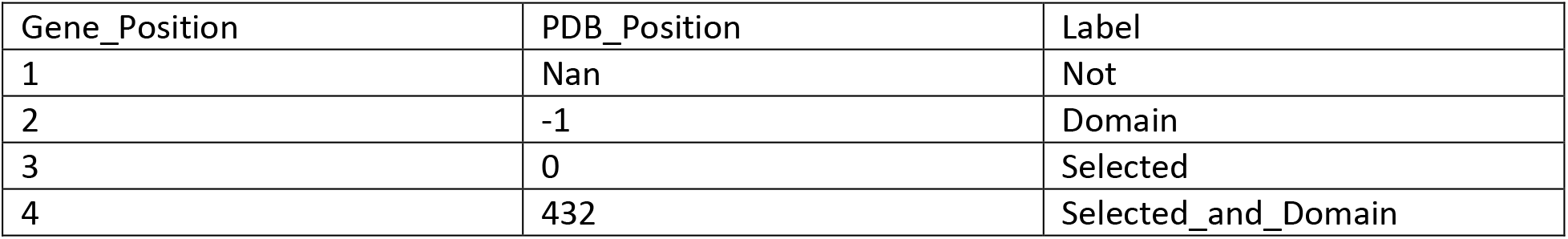
LYS table output of correspondence among the coordinates/residues of the studied sequences. These dataframes are directed to the Positions_Dataframe folder.

| Gene_Position | PDB_Position | Label |
| --- | --- | --- |
| 1 | Nan | Not |
| 2 | -1 | Domain |
| 3 | 0 | Selected |
| 4 | 432 | Selected_and_Domain |

*Main scripts*:

- LYS_PDB_Search.py: Performs a blast search against RSCB database to find and download the best PDB files for the query sequences. The results are saved to the files “Full_Blast_results_against_PDB.tsv” and the reduced version containing the best scoring results, “Full_Blast_results_against_PDB_Filtered.tsv”. This is followed by the generation of a data frame of the correspondent positions among each query sequence and the homologous sequence. Simultaneously these positions are assigned a label that indicates whether: a) “Domain” they belong to the domain residues (using Prosites [8,9]), “Selected” they are positively selected (given by the codeml [1] output), “Selected_and_Domain” both or “Not” none.
- LYS_PyMOL_input_Dataframe.py: Takes the output data frame of LYS_PDB_Search.py and generates a customizable graphic visualization.

*Complementary scripts*:

- LYS_PyMOL_Prosites.py: Inputs individual sequence and a chosen PDB file, and allows personalized configuration. The domain positions can be assigned using various methods, for example via Prosites [8], a list of “\n” separated positions (referring to the query sequence) or by using the desired Uniprot’s domain sequences clustered in a fasta file. They will be locally aligned to the PDB file sequence.
- LYS_PyMOL_GUI_Prosites.py: GUI version of LYS_PyMOL_Prosites.py

## 5. RESULTS

### Testing the Scripts

The scripts were tested in a Unix server on 5 protein coding sequences of 438, 244, 183, 122 and 61 amino acids long, which are available at https://github.com/LysSanzMoreta/LYS_Automatic_Search/tree/master/TestSequences, together with their corresponding codelm results. The LYS_PDB_Search.py script running time was measured and the results are 1m56.113s for real, 0m9.880s for user and 0m0.296s in sys times. These sequences contain several types of examples, such as some sequences that do not show homologous proteins, some only show one or several matches in the PDB database and one that contains positively selected residues that are present in the functional domain of the homologous protein (see Figure 3).

## 6. DISCUSSION

After detecting regions of the genome under fast evolution, one of the goals of molecular evolution studies is to understand the functional impact of mutations in those regions. It is already possible to pinpoint the positions in a certain protein that seem to be evolving at a fast rate, but to infer the impact of a mutation in the protein function *in silico* it is important to first map it to a protein structure, when available, or an adequate template corresponding to a homologous protein. LYS automates the search for protein structures, depicts them in PYMOL together with the information on known functional domains, and incorporates the information from PAML’s M8 output providing a publication-ready representation of the results. It also creates easy to parse tables with all the results, facilitating further analyses of the end user.

## ACKNOWLEDGMENTS

The authors gratefully acknowledge the following for supporting their research: Villum Fonden Young Investigator Grant VKR023446 (R.R.F. and L.S.M.); the Danish National Research Foundation for its support of the Center for Macroecology, Evolution, and Climate – grant DNRF96 (R.R.F.); Novo Nordisk Foundation grant NNF16OC0023494 (L.S.M.); Programa Operativo de Empleo Juvenil FSE 2104-2020 – grant CCI 2014ES05M9OP001 (L.S.M.).

## AUTHOR CONTRIBUTIONS

L.S.M. and R.R.F. designed the study; L.S.M. wrote the software with input from R.R.F.; L.S.M. wrote the manuscript with contributions from R.R.F.

